# Generative Neuromorphic Programming of Mammalian Cells

**DOI:** 10.64898/2026.09.18.752793

**Authors:** Jean Disset, Charles Van De Mark, Georg K. A. Wachter, Shuo-Hsiu Kuo, Andrew Moorman, Wenlong Xu, Eric Palanques Tost, Andres Buritica Monroy, Calin Belta, Ron Weiss

**Author notes:** These authors contributed equally to this work (Jean Disset, Charles Van De Mark).

## Abstract

Cells make decisions through densely interconnected regulatory networks that compute graded, analog responses[1, 2]. Synthetic biology has made cells programmable, but the field’s dominant abstractions poorly match the analog nature of cells. Designing analog circuits that work in cells is challenging: predicting how components combine remains difficult, and the vast combinatorial space of parts, weights and topologies exceeds what experiments can explore. Here we establish generative programming of mammalian cells through a neuromorphic framework that compiles desired behavior into DNA. Composable regulatory devices built from RNA-targeting endoribonucleases (ERNs) supply signed weights and nonlinear activation, enabling multi-layer circuits with diverse multi-input analog responses. To predict and design such circuits, we develop biomorphic neural networks (BMNs): a compositional architecture that mirrors biological interactions, in which each basic process (transcription, translation, cleavage) is captured by a reusable neural block learned from the behavior of whole circuits that contain it. Recomposed into architectures absent from training, these models predict responses with errors often comparable to variability between experimental repeats. Inverting this process, our software package, the biocompiler, jointly optimizes topology, parts, and weights to realize specified target behaviors. Challenged with three target behaviors, the biocompiler proposed three previously unseen architectures; we built all three, and each reproduced its target behavior in mammalian cells in a single design pass without manual tuning. Generative neuromorphic programming thus offers a systematic route to engineering the analog computation native to living cells.

## Introduction

Cells naturally make decisions by recognizing and processing multi-dimensional patterns of gene expression and cellular biomarkers[3]. This capacity emerges from densely interconnected signaling and gene regulatory networks that integrate many inputs to compute a response [4, 5, 6, 7]. Crucially, these input-output behaviors and computations are often analog: pathways such as NF-*κ*B, ERK/MAPK and p53 convert combinations of molecular signals into graded, dose- and context-dependent outputs, allowing a limited molecular repertoire to specify a vast range of cellular behaviors [1, 8, 9, 10, 11]. Programming similarly rich input-output functions is a long-standing goal of synthetic biology, and would unlock wide-ranging applications including cancer cell classification and destruction [12, 13, 14, 15], guided differentiation of organoids and tissues [16, 17, 18], cell therapy [19, 20], and biomanufacturing [21].

Yet, the tools we use to program cells remain mismatched to how cells natively compute. Synthetic gene circuit design is still dominated by Boolean abstractions [22], and existing computer-aided design workflows focus on digital logic [23]. Boolean circuits operate near or at saturation levels, compete for cellular resources, and can fail at scale[24]; new behaviors often require new topologies, and graded, analog responses can be difficult to represent compactly. One way to fundamentally solve this mismatch problem is to take another route: neuromorphic computing. Rather than imposing digital abstractions on an analog substrate, the neuromorphic paradigm exploits the substrate’s native nonlinear primitives [25]. Artificial Neural Networks (ANNs) illustrate this principle *in silico*, composing weighted, nonlinear elements into diverse complex functions. Because gene regulation is itself analog and compositional, cells are a natural substrate for neuromorphic computation and can provide suitable modular molecular primitives. What is needed is a design abstraction that allows one to wire and tune these primitives into circuits, together with a way to navigate the design space, and turn desired behavior into a genetic implementation.

Elements of neuromorphic computation have already been described in biological systems [26, 27, 28, 29, 30, 31], including *in vitro* DNA strand displacement [32, 33, 34], transcription-factor networks in bacteria [35, 36], and single-layer classifiers in mammalian cells, including a winner-take-all comparator [37] and a sequestration-based perceptron built from the same Csy4 endoribonuclease used here [38]. Signaling-based neural networks have also been studied theoretically [39]. Multi-layer circuits and gradient-based optimization have been demonstrated in bacteria[35]. These ingredients show that a general design framework for mammalian cells is within reach. Two important challenges remain. The first is to establish how neuromorphic primitives in mammalian cells can be composed into multi-layer circuits. The second is to turn a desired analog computation into a genetic program. Analog function is a graded surface rather than a truth table, component composition is nonlinear and difficult to anticipate, and the space of parts, weights and topologies grows far faster than experimental search can cover.

To this end, we establish generative programming of mammalian cells, realized through neuromorphic gene circuits (NGCs). We compose RNA-targeting endoribonucleases (ERNs) into networks, and build a design engine that can translate desired behavior into such networks and compile it into DNA. These genetic devices supply signed weights and nonlinear activation, enabling multi-layer NGCs that integrate signed inputs and pass graded outputs between nodes. We experimentally realize a broad repertoire of multi-input analog behaviors, spanning single-, dual- and triple-region responses, signaling cascades, and two-dimensional band-pass filters.

To predict and design such circuits, we introduce biomorphic neural networks (BMNs), differentiable models of the molecular processes underlying NGCs. Each process, such as transcription, translation, or RNA cleavage, is represented by a small neural network called a neural block, which learns the process’s quantitative input-output response rather than assuming a predetermined kinetic equation. A BMN is thus a network of networks: its neural blocks are connected according to the circuit’s molecular interactions, so a single ERN node is represented by several blocks. Blocks are learned and shared across circuits, and their behavior is never observed directly but rather inferred from composite circuit responses, so each new measurement constrains all processes reused across circuits, including for topologies absent from training. Exploiting this shared structure, our BMNs predicted unseen circuit responses with errors often comparable to variability between experimental repeats. Differentiability allows us to calculate how changes in design parameters affect the predicted response and use these sensitivities to guide NGC design. BMNs can also generate gene circuit designs for specified input-output responses. To leverage this, we built the biocompiler, a software suite that performs gradient-based design on BMNs, jointly optimizing topology, parts, and weights, to achieve a desired cellular response and convert it to DNA.

### Molecular computing primitives

At its core, a neural network composes a few elementary operations: each node computes a weighted sum of signed inputs, applies a nonlinear activation function, and passes its output downstream. ANNs realize these operations with signed real numbers *in silico*; here we leverage the native chemistry of cells to implement them in NGCs. We decompose these requirements into five primitives for which we find a corresponding biochemical implementation (Figure 1a): positive inputs through co-expression, negative inputs through active removal, weights through tunable genetic sequences and co-transfection ratios, nonlinear activation through ERN-mediated regulation, and composition by connecting one node’s output to the next. Negative inputs and nonlinear activation thus share a molecular mechanism.

**Figure 1.**
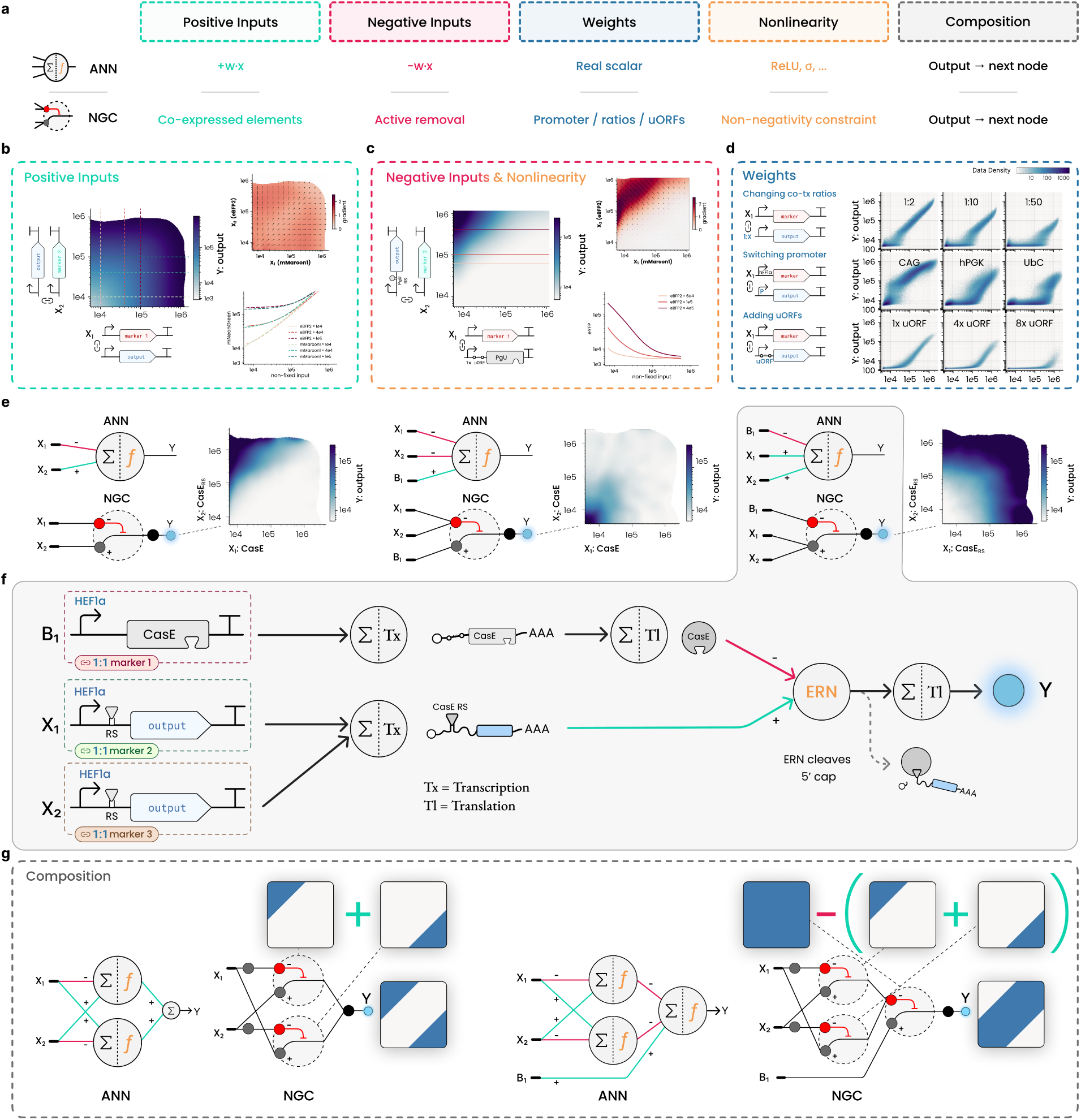
Primitives of neuromorphic computing in cells. **a**, The five primitives of neuromorphic computation (rows) and their counterparts in artificial neural networks (ANNs; left) and neuromorphic gene circuits (NGCs; right): positive inputs (weighted positive sum / co-expression of converging elements on a shared target mRNA), negative inputs (weighted negative sum / ERN-mediated removal of that mRNA), weights (scalars / promoter strength, 5’-UTR uORFs, co-transfection ratios), nonlinearity (activation / graded dose-response with basal floor and saturation), and composition (connecting one node’s output to another node’s input). **b**, Positive inputs. *Left*, 2D response with two co-expressed copies of the same output TU, each with its own marker; dashed lines mark the slices shown in 1D on the bottom right. *Top right*, local sensitivity on a log-log scale (arrows toward higher output), showing a relatively uniform (= quasi-linear) gradient distribution throughout the operating range. *Bottom right*, output vs positive input with three fixed levels per input axis levels (colour-matched). The lines match exactly, showing *X*_1_ and *X*_2_ marker independence. The lines show a near-perfect linear relationship (*y* = *X*_1_ + *X*_2_, where one input is fixed). **c**, Negative inputs and nonlinearity (PgU ERN node). *Left*, 2D response with mRNA input on *X*_2_ and ERN input on *X*_1_; dashed lines mark the three slices on the bottom right. *Top right*, local sensitivity on a log-log scale (arrows toward higher output), showing a dim basal floor at high ERN, where the mRNA is already cleared, and a brighter region where either input still moves the output. *Bottom right*, output along the positive input at three fixed ERN levels (colour-matched); the gap from each linear reference (fit through the same endpoints) is the node’s nonlinearity, whose shape the BMN learns from data. **d**, Weight tunability on a single output TU. Top: co-transfection ratios (1:2, 1:10, 1:50) give continuous modulation. Middle: promoters of different strengths (CAG, hPGK, UbC) set expression at defined levels. Bottom: repeated 5’-UTR uORFs (1×, 4×, 8×) attenuate across nearly two orders of magnitude. **e**, Three single-neuron examples, each shown as an ANN diagram (top left), its equivalent NGC schematic (below), and the experimental 2D dose-response over cotransfection markers *X*_1_, *X*_2_ (right): one positive and one negative input (left); positive bias with two inputs on the negative (ERN) hub (middle); negative bias with two inputs on the positive (mRNA) hub (right). **f**, Molecular zoom of the rightmost example in **e**: the negative-side TU encodes the ERN protein, which cleaves matching sites on the output mRNA, removing it from the shared reaction pool and producing the node’s nonlinear response. **g**, Composition: a single-layer dual-region pattern (left) and a two-layer 2D bandpass (right). Throughout the manuscript, for 2D dose-response maps, axes are the molecular inputs, colour is the node or circuit output, and the input space is sampled by poly-co-transfection (further described in Methods).

A major difference between ANNs and NGCs comes from ANNs combining the sign and magnitude of a weight into a single signed scalar *w_i_*. Because a concentration cannot carry a sign, NGCs separate these two properties. The sign is set by how an input acts: an input can contribute positively, by adding to a node’s output, or negatively, by removing it. Positive contributions arise from co-expression of molecular elements that converge on a shared output, e.g. different DNA constructs that produce the same mRNA, or distinct mRNAs that translate into the same protein. We can implement a simple node with two positive inputs by using two copies of the same transcription unit (TU), each tracked by an independent co-transfection marker. Throughout, we use poly-co-transfection [40] to deliver a different combination of input levels to each cell in a single sample, so that the cell population collectively samples the entire input space in one experiment (Methods). Experimentally, the output combines the two inputs additively to a very good approximation, with gradient magnitudes of 0.8–1.0 across the input space (Figure 1b).

The negative contribution and the nonlinearity can both be supplied by an ERN, making it the core element of NGCs. Each ERN binds its target site on the output mRNA and triggers degradation via 5’-cap removal [41], lowering the steady-state concentration of surviving mRNA. Unlike an artificial neuron, which combines signed inputs before applying a separate activation function, the ERN node combines supply, removal, and nonlinear activation in the same physical process. Output remains near a basal floor where removal dominates, and rises as positive supply increases. This graded non-linear transition constitutes our activation function. The nonlinearity is directly visible in the two-dimensional dose-response of a PgU node (Figure 1c), with flat and positively sloped regions. We do not prescribe a kinetic form for this transition: its precise shape is learned from data by the BMN. We use three orthogonal ERNs (PgU, CasE and Csy4, drawn from the Cas6 and Cas13 families [41]), whose distinct cleavage specificities and RNA-removal strengths enable independent connections; each ERN’s characteristic activation function provides a further tuning parameter.

Weight magnitudes set the relative strength of each input. Here we implement weights via three mechanisms: varying co-transfection ratios, using promoters of different strengths, and inserting upstream open reading frames (uORFs) into the 5’ UTR[40]. Each offers a distinct lever on the weight of an input: co-transfection ratios provide precise, continuous modulation (Figure 1d, top); promoters allow to tune transcriptional expression, (Figure 1d, middle), and uORFs give graded post-transcriptional attenuation across nearly two orders of magnitude (Figure 1d, bottom; and demonstrated previously[40]).

Figure 1e presents three multi-input single-node examples as ANN diagrams, equivalent NGCs, and measured two-dimensional ⟨*x*_1_, *x*_2_⟩ dose-responses. Our NGC notation mirrors ANN diagrams while reflecting the underlying biomolecular processes. Figure 1f details the rightmost node; the others use the same machinery and differ only in which hub each input feeds. Each ERN node has two input hubs: a positive hub (transcription producing output mRNA) and a negative hub (translation producing ERN protein). Wiring inputs to either hub allows arbitrary combinations of positive and negative contributions. Either hub can also receive a fixed input, a *bias*, analogous to an artificial neuron’s constant term[42]. The direct output of an ERN node is therefore the surviving amount of output mRNA.

Nodes can be combined by having their surviving mRNAs encode the same reporter, so that translation of these mRNAs contributes to a shared protein output. The surviving mRNA output can also serve as an input to any node in a subsequent layer, allowing nodes to be composed into multi-layer circuits. For an inhibitory connection, this mRNA encodes an ERN that cleaves the downstream node’s target mRNA. Translation therefore converts the upstream RNA output into a downstream inhibitory input: RNA survival in one layer controls RNA removal in the next. Circuit inputs can also be wired directly to any node in any layer, contributing to target-mRNA or ERN production.

An example one-layer, two-node network has two nodes producing the same reporter in opposite corners of the input space; their summed outputs form a dual-region pattern. An example two-layer, three-node network replicates the structure of the first example but feeds the output into the negative hub of the second-layer node biased toward constitutive high expression which yields a diagonal two-dimensional bandpass (Figure 1g).

### Learning and designing circuits

The primitives of neuromorphic computation compose into a combinatorially vast space of parts, weights and topologies, far too large for exhaustive experimental characterization. To predict and design circuits across this space, the biocompiler trains a single underlying model on a corpus of circuits, representing each as a biomorphic neural network (BMN), and applies it in two directions: prediction, which maps a specified gene circuit to its function, and design, which maps a desired function to a proposed circuit. Because biological operations recur across NGCs, and BMNs capture that reuse, the learned model generalizes in both modes to combinations of parts, weights and topologies absent from training.

Figure 2a illustrates the conversion between two representations of an NGC: a genetic recipe (genetic parts and transfection protocol) and a BMN compute graph, representing molecular processes the recipe gives rise to in the cell. Each node in this graph is a biological process (e.g., transcription, translation, ERN cleavage), and each edge a molecular species produced or consumed by the processes it connects. Constructs delivered together feed into a common node parameterized by their mixing ratios, allowing design to adjust those ratios by gradient descent. The conversion runs in both directions. The forward map, used in training and prediction, applies biological graph-rewriting rules: DNA is transcribed into RNA, RNA is translated into protein, and ERNs cleave their target transcripts. Each rule instantiates the corresponding nodes and edges. The reverse map underlies design, running the same rules backward to generate a genetic recipe from a compute graph.

**Figure 2.**
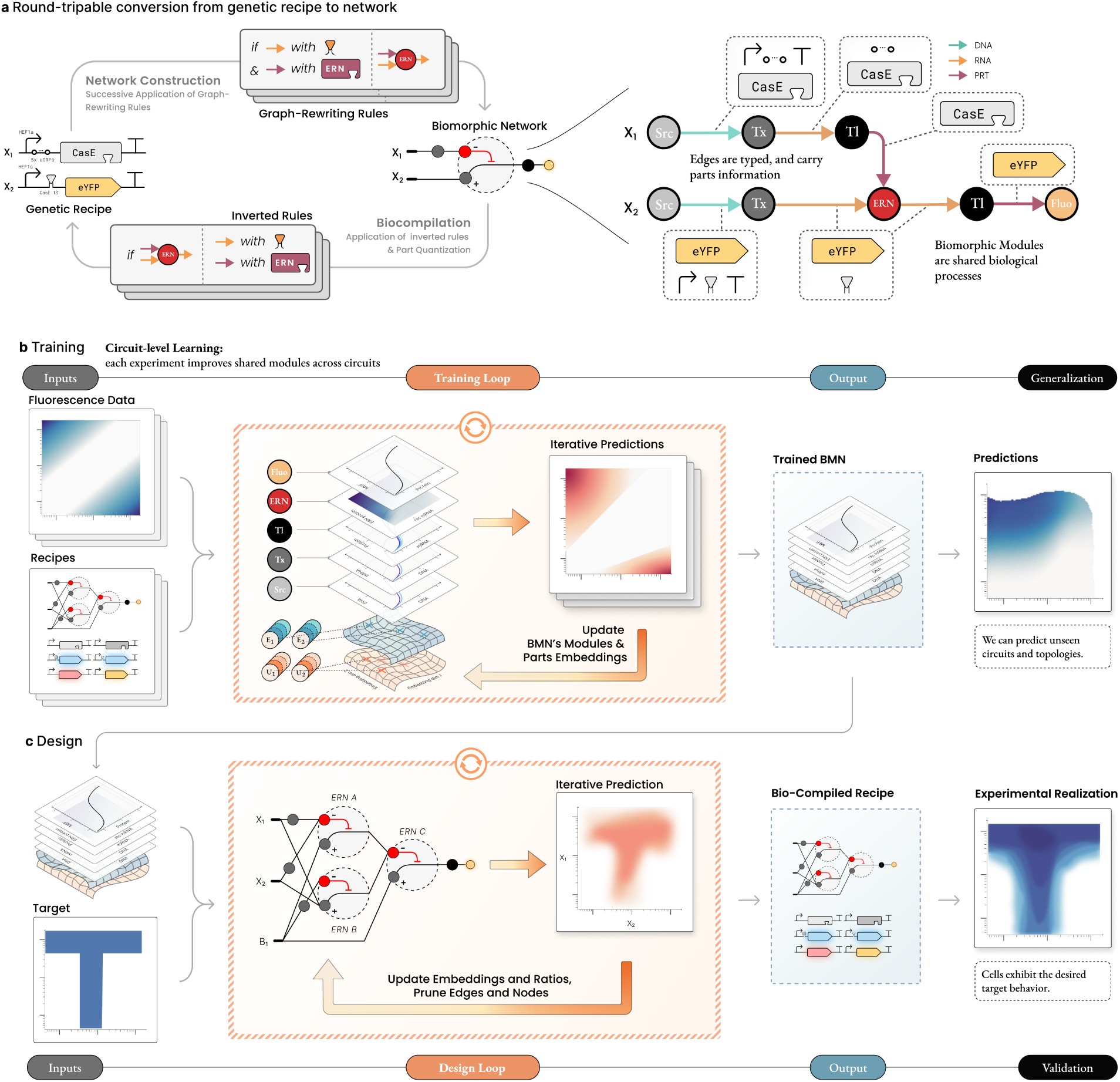
Biomorphic neural network (BMN) architecture: from genetic recipe (DNA construct specification and transfection protocol) to compute graph, training, and design. **a**, Recipe-to-network conversion. A sequence of graph-rewriting rules turns each genetic recipe into a circuit-specific biomorphic network (network construction); the same rules applied in reverse, together with part quantization, compile a BMN back into a buildable recipe (biocompilation). The same graph-rewriting machinery runs in both directions, so it serves both prediction and design. The resulting biomorphic network is a typed compute graph: edges are typed by molecular species (DNA, RNA, protein) and carry the embeddings of the parts they contain, and nodes (Src, Tx, Tl, ERN, Fluo) are shared blocks reused across all circuits. **b**, Training mode. Fluorescence data and genetic recipes are translated into compute graphs, and a training loop iteratively updates the shared blocks and part embeddings until predictions match observations across all circuits. The trained BMN generalizes to unseen circuit architectures and topologies (right). **c**, Design mode. The process runs in reverse: given a target cellular response (one of the target patterns shown in Figure 6b), the biocompiler initializes a densely connected scaffold graph and iteratively updates embeddings and transfection ratios while pruning edges and transcription units until the prediction matches the target. The output is a bio-compiled genetic recipe whose experimental realization recapitulates the desired behavior.

In a BMN’s compute graph, each node is itself a small neural network, a *neural block*, that predicts the output of one biological process from its inputs, and these blocks are learned from training data rather than specified in advance. Representing each regulatory process this way, rather than as a mechanistic rate law or a hand-tuned parametric function, lets a BMN capture the input-output behavior these processes actually exhibit, which is typically nonlinear, saturating, and dependent on how much of each input is present, to the level of accuracy the data support. It does so without committing to a mechanistic form or to kinetic parameters that, for most genetic parts in our library, have never been measured directly in a cellular context. The cell type a circuit runs in is supplied to every block the same way, as an embedding learned from data. With every node encoded as a block and edges carrying the molecular species that pass between them, each BMN can itself be treated as a single neural network. Crucially, training a BMN uses measurements of whole-circuit behavior; the individual blocks are never observed directly. Because the same block recurs across many circuits, every measured circuit that contains it constrains that block’s behavior, so a transcription, translation, or cleavage block is learned jointly from the entire corpus of circuits in which it appears. Any experiment expressible within the recipe-to-graph framework can therefore contribute to these shared models: knowledge accumulates across designs rather than being confined to a fit for each circuit. This reuse also lets the model generalize to circuits it has never seen.

Figure 2b presents an overview of this training phase, which precedes any prediction or design of new circuits. We train on a set of genetic recipes together with the fluorescence data measured in the corresponding transfection experiments. Each recipe is first converted into its BMN, then the biocompiler trains all of these networks simultaneously, optimizing the parameters of the shared blocks so that every network reproduces its observed fluorescence data. Once trained, these blocks are available to be recomposed into any circuit the biocompiler needs to predict or design, including graph topologies absent from training.

Figure 2c shows the design process, which uses the trained blocks to solve the inverse problem: rather than fitting blocks to observed data, the biocompiler holds their learned behavior fixed and searches for circuits that produce a target input–output pattern. Genetic parts are naturally discrete, not a continuous dial that gradients can search directly. To bridge this gap, each experimentally available part is assigned a position in a continuous *latent space*, an *embedding* that summarizes its effect on circuit behavior. These positions are learned jointly with the blocks from the same circuit data during training, placing parts with similar effects close together. Gradient descent can then optimize a continuous position before quantizing it to the nearest real part. Starting from a candidate scaffold, gradient descent adjusts co-transfection ratios and part choices; differentiable masks additionally allow connections and transcription units to be pruned (Methods). Separate runs can settle on different candidate solutions.

The result of this search and optimization process can finally be compiled into a complete, buildable genetic recipe that realizes the target behavior as closely as possible.

The processes introduced in this section (BMN architecture, training, and design procedures) are described in detail in the Methods.

### Circuit composition and predictions

We designed our next experiment to demonstrate two things: that uORF-based weights generate a customizable range of analog input-output functions, and that shared BMN blocks predict new part combinations from characterization of just a few. Nine uORF variants on both target and ERN sides define an 81-condition tuning matrix for a single PgU node (Figure 3a). Increasing uORF-mediated attenuation on the ERN side weakens the negative input, expanding the high-output region toward the bottom-right. Increasing attenuation on the fluorescent-protein side instead shifts this region toward the top-left and decreases signal strength. To the left of the matrix, nine uORF:EYFP measurements independently characterize the variants’ effects on translation.

**Figure 3.**
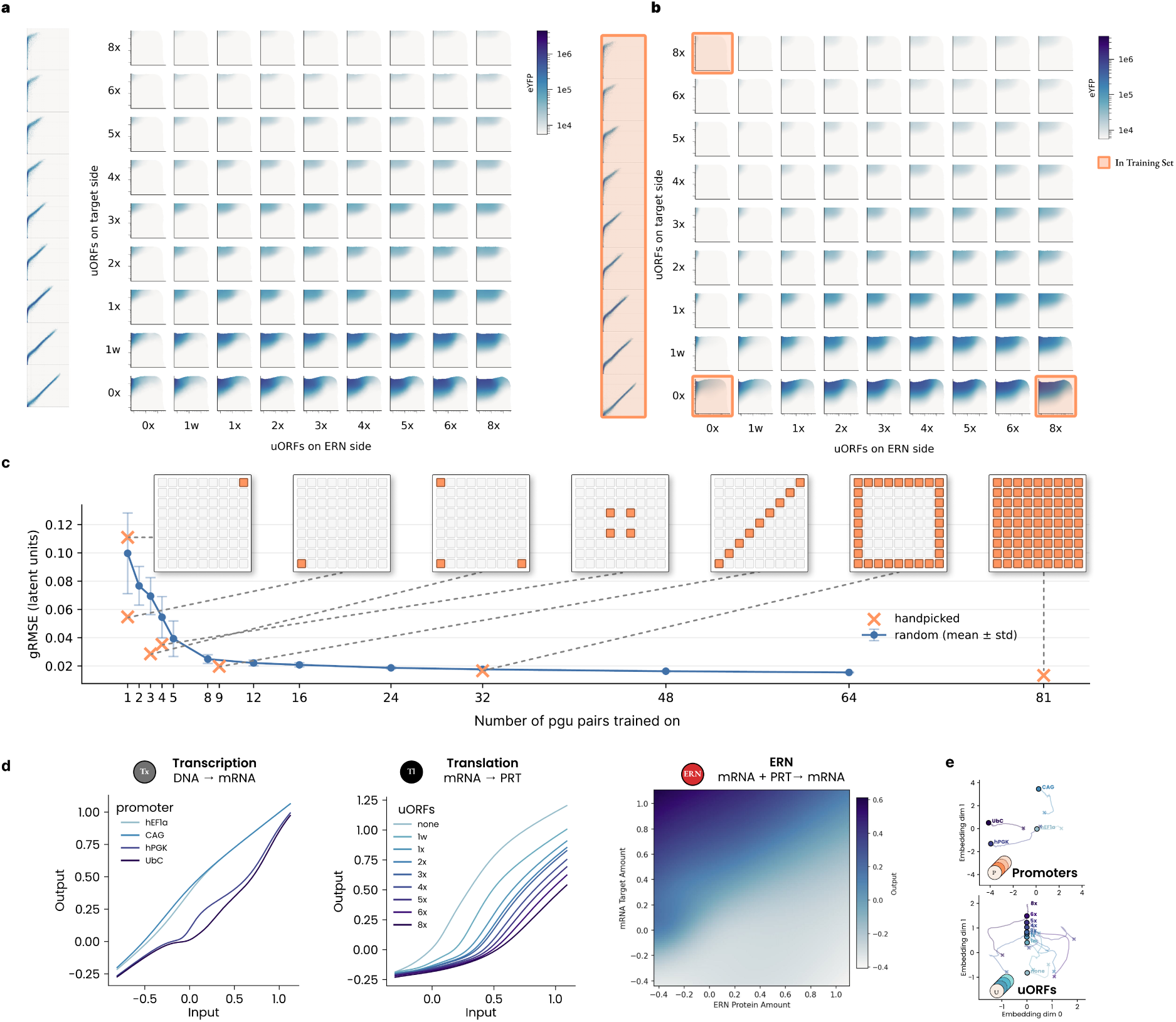
Circuit-level learning and generalization on the uORF tuning matrix. **a**, Experimental single-node data. Left column: fluorescence of eYFP with 0 to 8 uORFs versus an eBFP2 reference. Matrix: PgU-based ERN-node output across 81 uORF combinations on the target mRNA side (y-axis, positive input) and ERN-encoding side (x-axis, negative input), showing systematic shifts in the transition between low and high output. **b**, Biocompiler predictions for the full matrix. Orange-outlined squares indicate the training set (3 of 81 PgU-based ERN-node conditions plus nine uORF:EYFP transcription units). The remaining 78 conditions are predicted from leveraging the trained shared blocks and uORF embeddings. **c**, Prediction performance scaling. gRMSE (rescaled units, 0-1) versus training set size (k out of 81 conditions plus nine uORF transcription units). Circles are the mean over 30 randomly drawn subsets at each k, each trained from scratch, and bars are one standard deviation over those 30 models. Insets show specific cases for several values of k with matrix positions indicated. Carefully picked selections (orange crosses) can approach larger training sets when they span the matrix (e.g., 3 corners gets most of the way to the 9-element diagonal). Each model is evaluated over all 81 matrix conditions, including those it was trained on; restricting the evaluation to held-out conditions only changes the curve by less than 0.001 gRMSE. **d**, Learned blocks after training with the nine uORF:EYFP TUs, full PgU matrix, and four promoters constitutively expressing EYFP. From left to right: transcription (Tx), showing DNA-to-mRNA mapping for four promoters tested; translation (Tl), showing uORF-dependent nonlinear modulation of mRNA-to-protein conversion (curves colored by uORF count); ERN (PgU), showing the learned RNA removal function: the target mRNA surviving cleavage as ERN protein (X axis) and target mRNA levels (Y axis) vary. **e**, promoter and uORF embedding trajectories during a training run, showing the learned latent-space positions of the four promoters (top) and uORF parts (bottom; 0x to 8x) arranged along a smooth gradient, with the optimization path from random initialization (crosses) to final positions (circles).

Full characterization of all uORF combinations would require 81 individual experiments for a single ERN node. Here we show that we can learn to extrapolate (and interpolate) uORF behavior on both sides of the ERN node by training on basic uORF experiments and very few instances of the full matrix. Figure 3a shows the experimental ground truth, and Figure 3b shows the predictions produced by a BMN trained only on the basic uORF set plus the 3 orange corners of the matrix. The held-out matrix elements are predicted by reusing the same learned transcription, translation and ERN blocks across unseen uORF combinations.

Throughout this work, prediction error is measured by the root mean square error across the input-space grid (gRMSE) in rescaled units (0-1). When biological repeats are available for a given circuit, we can compute two directly comparable metrics: pred gRMSE is the gRMSE of a prediction by a BMN model, and rep gRMSE is the average gRMSE between two or more repeat experiments of the same circuit. The prediction over the entire PgU matrix in Figure 3b reaches a pred gRMSE of 0.02, compared with a rep gRMSE of about 0.05 across 24 separately characterized circuits measured in two or more independent experiments.

Figure 3c shows the effect of training set size on prediction accuracy. As the number of training conditions increases, pred gRMSE decreases with rapidly diminishing returns. Notably, the selection of training conditions matters as much as their number: picking three corners of the matrix (pred gRMSE=0.029) gets most of the way to an entire diagonal of nine conditions (pred gRMSE=0.020). Increasing the training set to the full bounding box (32 conditions) only marginally improves pred gRMSE to 0.017. We performed the same prediction workflow on the remaining ERNs, yielding similar results.

Figure 3d shows the learned neural blocks after training a BMN model on the full PgU matrix, the uORF:EYFP transcription units, and our four promoters, of which only hEF1a was used in the matrix and uORF experiments. The transcription block (Tx) shows the DNA-to-mRNA mapping of the four promoters characterized for this training set, representing an inferred relationship between gene copy number and transcript levels. The translation block (Tl) reveals uORF-dependent nonlinear modulation as increasing uORF count progressively attenuates translation output. The ERN block (PgU) learns a two-input RNA-removal function capturing how ERN abundance reduces target mRNA. These are effective functions inferred from whole-circuit measurements, not independently measured molecular rates. Figure 3e shows the promoter and uORF embedding trajectories during training. The uORF embedding trajectory shows that the discrete uORF parts self-organize during training into a smooth, ordered gradient in latent space, correctly recovering their order by uORF count. This continuous ordering matters because gradient-based design requires a differentiable space in which to perform optimization steps, but genetic parts themselves are discrete and cannot be searched this way directly. By learning a continuous embedding in which discrete parts fall along a meaningful order, the biocompiler can instead search this analog space continuously through most of the design process, then quantize the final result to the nearest available part.

Having shown that the biocompiler generalizes to unseen weight combinations within a single node topology, we now scale up to multi-node circuits to test the expressivity that composability unlocks. Composability, the ability to chain one neuron’s output into another’s input, is a defining requirement of neuromorphic computing: it is what allows a small set of primitives to be combined into arbitrarily complex functions. To demonstrate this versatility experimentally, we built NGCs of increasing complexity from the same molecular nodes used above. This then allowed us to investigate whether the BMN architecture generalizes across topologies as well as weights, testing the biocompiler’s ability to predict architectures absent from its training data.

We built several single-layer compositions, showing that a small number of ERN nodes produce diverse analog functions from weight, bias and wiring changes alone. We created two-node networks using CasE and PgU that partition the input space into dual regions of high expression along opposing diagonals (Figure 4a). A first variant wires one input as positive to one node and negative to the other, and the second input in the reverse configuration, producing top-left and bottom-right high-expression regions with dynamic ranges exceeding 30-fold. A second variant rewires both inputs onto the positive hub of one node and the negative hub of the other, adding a constitutive bias to the remaining hub of each node, shifting high expression to the bottom-left and top-right. For each of these, the biocompiler, which saw no single-layer multi-node circuit during training, correctly predicts the input-output pattern of both circuits, including the reduced dynamic range of the bottom-left/top-right variant. We then extended to three nodes (PgU, CasE and Csy4), generating triple-region patterns in both 2D and 3D (Figure 4b): two-input circuits (with constitutive bias) produce three adjacent high-expression regions (top-left, bottom-right and top-right), while three-input circuits yield three distinct corners in 3D space. A BMN model that has never seen these 3-ERN circuits predicts their experimental behavior with accuracy comparable to differences between biological repeats.

**Figure 4.**
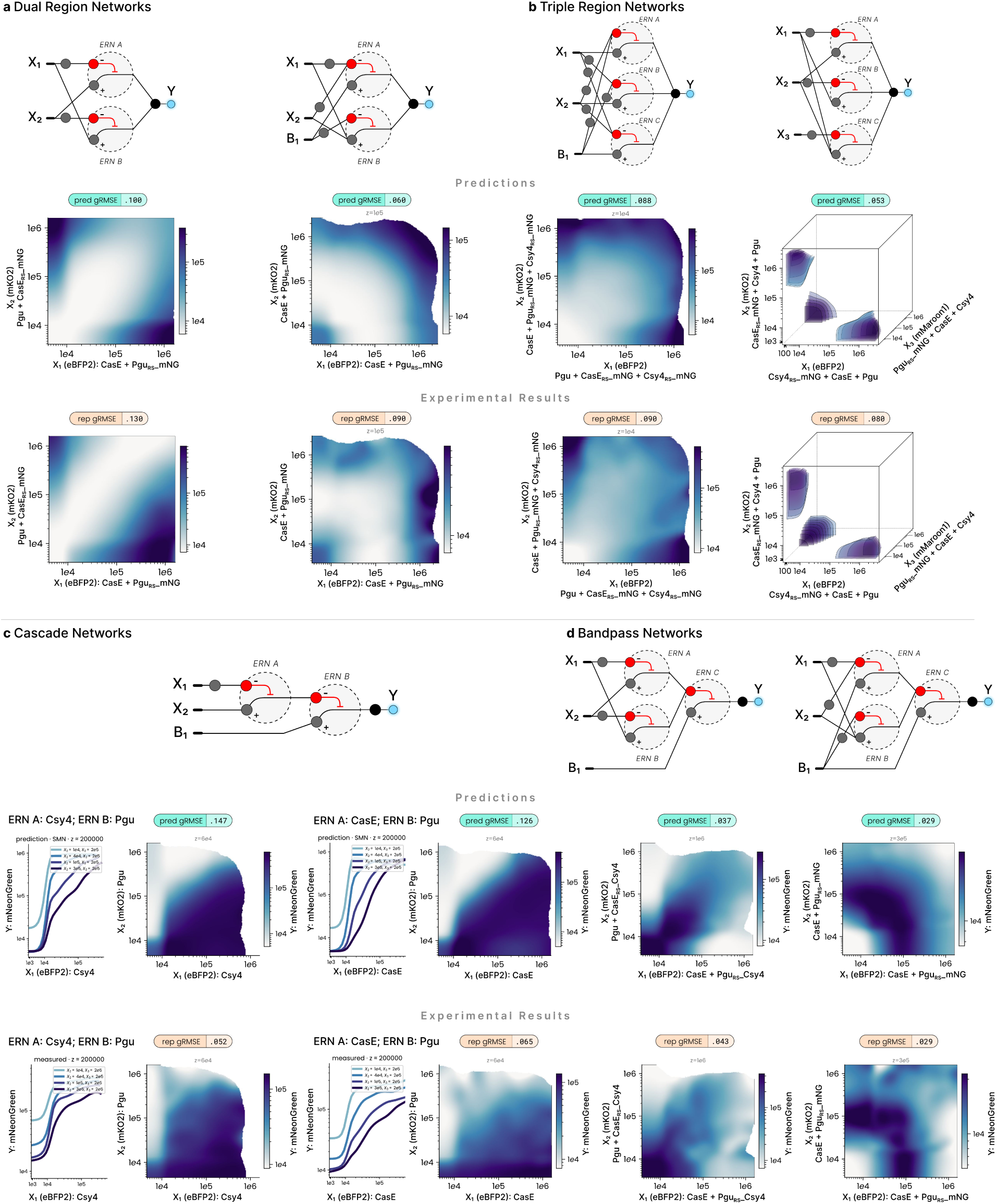
Multi-node multi-layer neuromorphic networks. **a**, Dual region networks. Two single-layer, 2-node architectures partition the input space into two high-expression regions along opposing diagonals. Left: top-left/bottom-right regions via complementary inhibition. Right: bottom-left/top-right regions. Predictions and experimental results are shown below each network schematic. Biocompiler predictions and repeat gRMSEs quantify how each prediction/repeat compares to another experiment; predictions are close to experimental noise. **b**, Triple region networks. Two 3-node, single-layer architectures extend partitioning to three distinct expression regions. Left: a two-input circuit with bias produces three adjacent regions in 2D. Right: a three-input circuit creates three corners of high expression in 3D input space. **c**, Cascade networks. A two-layer serial architecture where the output of the first ERN node feeds as input to the second. Line-plot view of slices are shown at different X_2_ levels. **d**, Bandpass networks. Two three-node, two-layer architectures produce 2D band-shaped regions of high expression. Left: bottom-left to top-right orientation. Right: top-left to bottom-right orientation. For all circuits above, the predicted topology class is absent from training. Dual- and triple-region variants share a model trained without multi-node circuits; cascades are predicted from single-layer data. The left bandpass model excludes left bandpasses; the right bandpass model excludes both bandpass classes.

As with ANNs, we can compose NGCs depth-wise into layers. In our two-node, two-layer cascade architectures (Figure 4c), the output of the first node feeds as a negative input to the second, with a bias serving as a positive input to this second node. Swapping which ERN occupies each layer changes both the output magnitude and the steepness of the resulting function. A BMN trained only on single-layer data recovers cascade behavior qualitatively, with larger quantitative errors than for the other topologies (see below). In Figure 4d, we show experimentally that three-node, two-layer networks yield 2D bandpass behaviors, using the theoretical NGC design introduced in Figure 1g. Here, the last-layer node sums the outputs of the first two nodes, which together form a dual region pattern, and subtracts this sum from a positive input bias, yielding a bottom-left to top-right bandpass. This bandpass achieves a *>*15-fold dynamic range. By rewiring the topology so that both inputs converge on the negative hub of one first-layer node and the positive hub of the other, with a bias connecting to each node’s other input hub, the resulting network instead produces a top-left to bottom-right bandpass with approximately 10-fold dynamic range.

Across all compositions, we evaluated a BMN model that has never seen the topology class it is asked to predict. The model recovers the arrangement of low and high-expression regions across these architectures, although dynamic ranges are not always captured perfectly. Prediction gRMSE is at or below repeat gRMSE for six of the eight compositions; the two serial cascades have larger quantitative errors but still qualitatively mostly recover the expected behavior. The complete set of held-out predictions and experimental data, spanning all topology classes and training sets, is shown in the Supplementary Information.

### Generalization across circuits and cells

We next examine how a model’s prediction accuracy depends on the circuit topologies included in its training, and whether behavior learned in one cell type predicts circuit performance in another. We first observe that the BMN blocks learned from training on the complete corpus of circuits (Figure 5a) are qualitatively similar to, but quantitatively distinct from, those learned from the uORF matrix alone (Figure 3d). Motivated by this difference, we then examined how the circuit classes used for training affect prediction accuracy across held-out classes. To analyze this cross-class transfer, we split the dataset into five classes: circuits with one ERN node or none (S), multi-node single-layer circuits (M), two-node cascades (Lc), left three-node bandpass circuits (Lbl), and right three-node bandpass circuits (Lbr). We then trained a separate BMN on every combination of these classes and measured its prediction accuracy (gRMSE) on each class absent from that training combination. Figure 5b presents these results as a matrix of simplified average effect on gRMSE values, one per training-class/target-class pair, quantifying how much each training class contributes to prediction accuracy on the corresponding target class. Overall, single-layer training data strongly contributes to single-layer predictions and multi-layer training data strongly contributes to multi-layer predictions, with single-layer training contributing somewhat more to multi-layer predictions than multi-layer training contributes to single-layer predictions.

**Figure 5.**
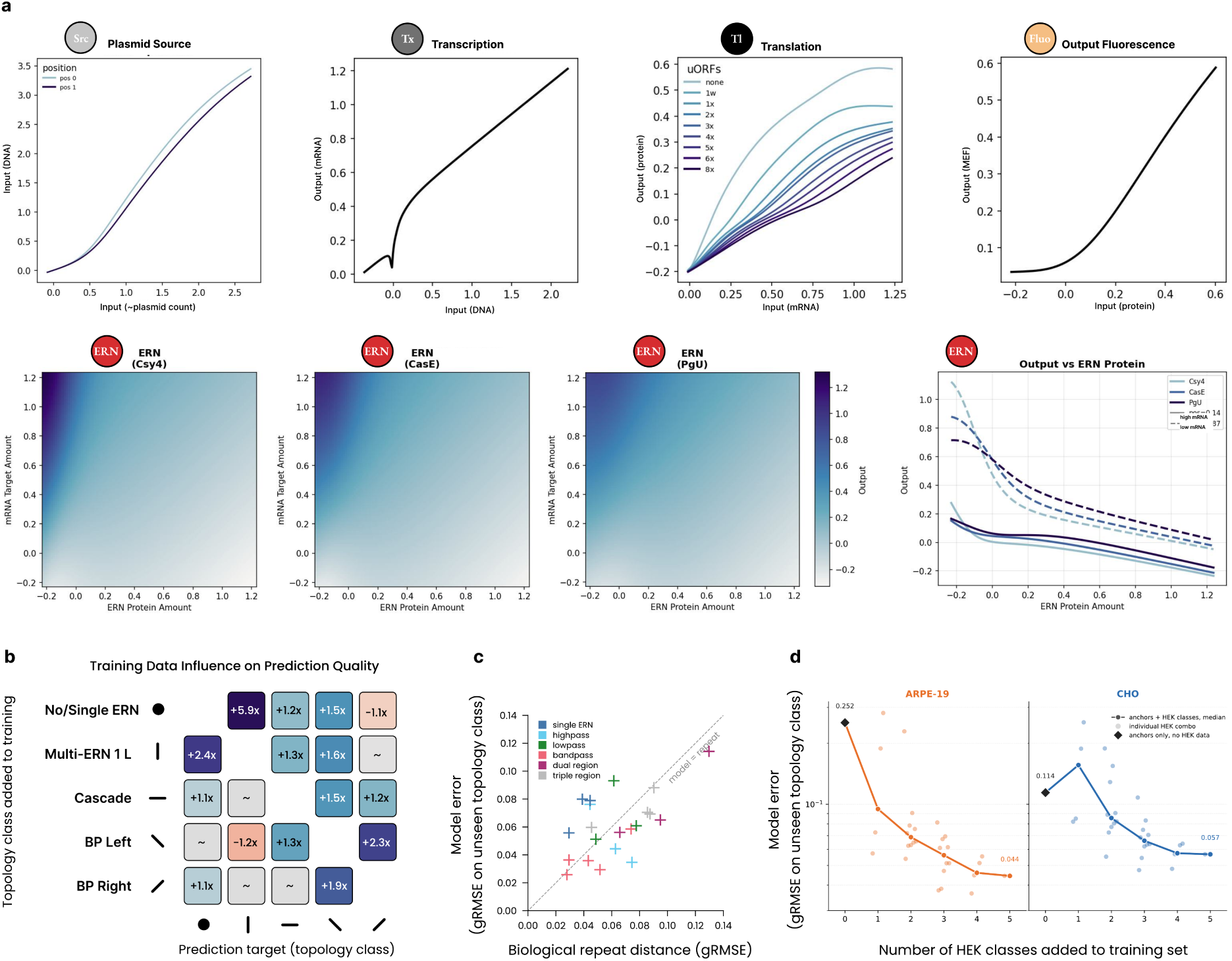
Learned blocks and generalization across topologies and cell types. **a**, The learned blocks after training on the full circuit corpus. Top row: plasmid source (DNA level as a function of plasmid copy number, shown for two transcription-unit positions), transcription (mRNA versus DNA), translation (protein versus mRNA, one curve per uORF count from none to 8×), and output-fluorescence (MEF versus protein) blocks. Bottom row: the three ERN blocks (Csy4, CasE, PgU) as two-input dose-response surfaces over ERN-protein and target-mRNA abundance, and their output along the ERN-protein axis at low and high target-mRNA levels (solid and dashed, respectively). Each block is shared: the same transcription, translation or ERN block is reused in every circuit where that process occurs. **b**, Cross-class transfer. For each pair of topology classes, the change in prediction quality on a target class (column) when a training class (row) is added to the training set, as a fold change (blue, improvement; peach, slight degradation; ~, negligible). Classes: no/single-ERN nodes, multi-ERN single-layer, cascades, and left and right three-node bandpasses. **c**, Prediction error versus reproducibility. For each held-out topology, the model’s error on the unseen class (prediction gRMSE) is plotted versus the distance between biological repeats of the same circuits (repeat gRMSE), coloured by circuit type. **d**, Cross-cell-line generalization. Model error on held-out topology classes in ARPE-19 (orange) and CHO (blue) cells as HEK293 topology classes are added to the training set. Black diamond, anchors only (the six single-ERN circuits measured in the target line) with no HEK293 data; light points, individual HEK293 class combinations; line, median.

To get a better sense of the overall performance of these held-out predictions, we built Figure 5c, which compares the model’s prediction gRMSE to the repeat gRMSE. Prediction errors cluster near the diagonal, and are therefore most often comparable to differences between experimental repeats, with a few topology-dependent departures from the line of equality. These results together suggest that the BMN is able to learn generalizable behavior from a subset of circuit topologies, and that the learned behavior is largely transferable to held-out topologies.

We next tested whether a model trained in one cell line could predict circuit responses in other lines without using their measurements for training. We built twelve circuits chosen to span all topology classes and measured them in ARPE-19, CHO and HEK293 cells. Circuit behavior was largely conserved across these lines, and a BMN trained only on HEK293 predicted the other two lines about as well as HEK293 itself (median gRMSE 0.031 in ARPE-19 and 0.044 in CHO, against 0.045 in HEK293). This suggests that the BMN is able to learn generalizable behavior across cell types, and that the learned behavior is largely transferable to held-out cell types.

We then asked whether HEK293 measurements could improve predictions when a small characterization set was already available in the target cell line. For each target line, ARPE-19 or CHO, we trained models using six single-ERN circuits measured in that line and added HEK293 data from different combinations of topology classes. As more HEK293 classes were included, median prediction gRMSE decreased (Figure 5d). Thus, in addition to predicting responses without target-line training data, the shared process models allowed HEK293 measurements to improve predictions when limited target-line characterization was available.

### Model inversion designs new circuits

Finally, one of the core motivations for this work is to enable the design of neuromorphic gene circuits that produce a desired cellular behavior. Having tested their predictive accuracy, we now use trained BMNs as generative models that map a target input-output function directly onto biological implementations. We chose three target patterns over two inputs, none of them resembling anything in the training data, so that the model had no familiar shape to rely on (Figure 6b). For each target, specified as a greyscale input-output map, the biocompiler searched within supplied generic scaffold architectures and a library of parts it had seen in whole-circuit measurements during training. Gradient descent adjusted part embeddings, transfection ratios and topologies (through pruning) to produce a buildable recipe without manual tuning of the generated design (Figure 6a). During the final step of biocompilation (conversion of the final compute graph into a genetic recipe), the biocompiler automatically quantized the optimized part embeddings to real, available parts, so that the optimization’s output was a complete DNA construct specification and transfection protocol (Figure 6f). The biocompiler returned three distinct circuit architectures (Figure 6c), none of which appeared in the training data (two dual-node circuits and one single-node circuit).

**Figure 6.**
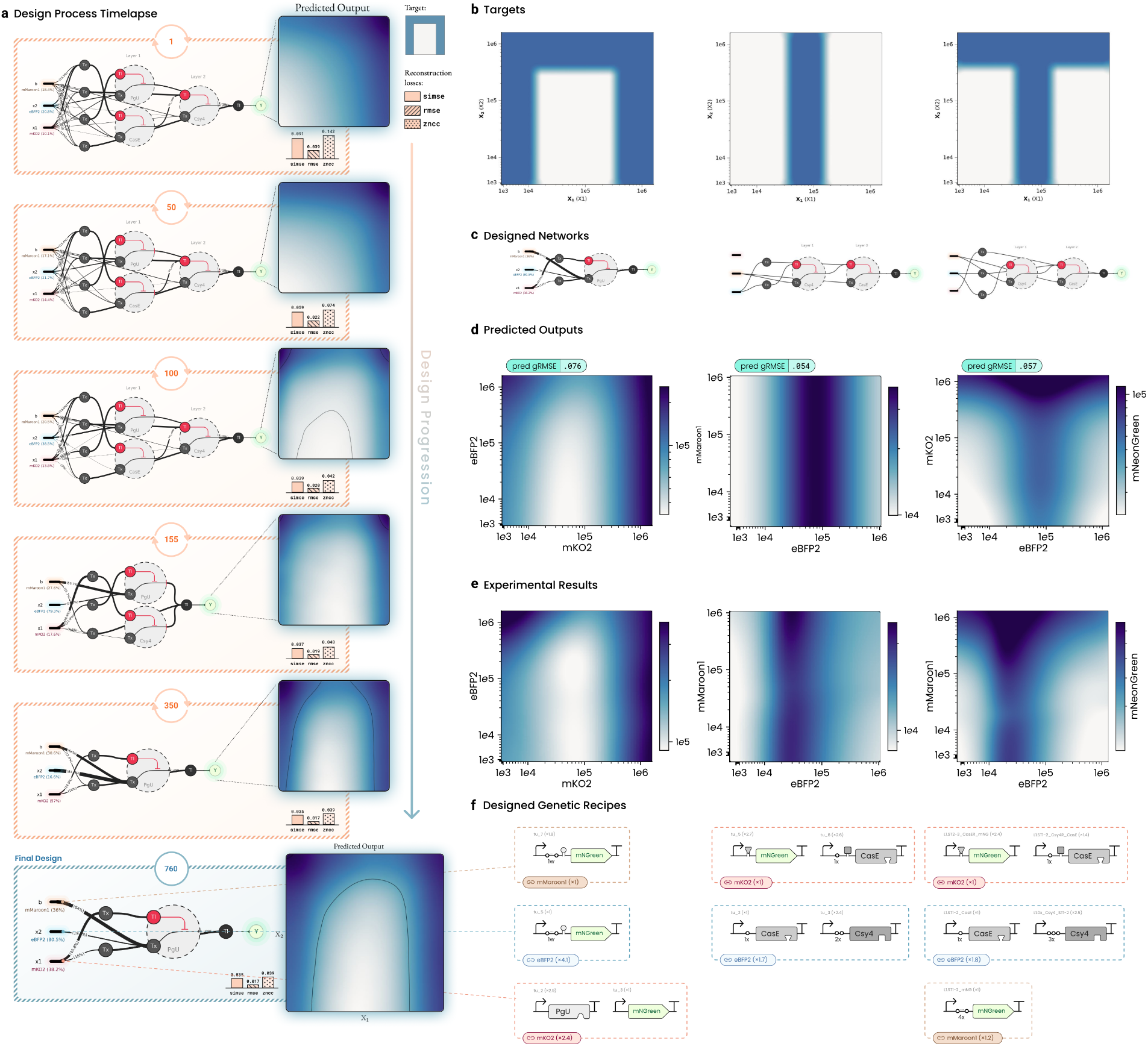
Generative design: from target patterns to experimental validation. **a**, Design process timelapse for one target pattern. Starting from a densely connected two-layer scaffold (top), the biocompiler iteratively updates part embeddings and transfection ratios while pruning edges and transcription units (dashed lines indicate pruned elements). Insets show the predicted output and reconstruction losses (SIMSE, RMSE, ZNCC) at each stage. Over 760 optimization steps, the scaffold is progressively simplified into a compact circuit architecture. Bottom: the final designed network with assigned parts and the corresponding genetic recipe showing DNA constructs and transfection ratios. **b**, Three user-specified target patterns provided to the biocompiler over two input concentrations (X_1_, X_2_). **c**, Circuit architectures designed by the biocompiler for each target. Each was pruned from a densely connected scaffold to a distinct architecture through gradient-based optimization. **d**, Predicted outputs of the designed circuits. **e**, Experimental measurements of the designed circuits after construction and transfection, recapitulating the target patterns. **f**, Designed genetic recipes for each circuit, showing DNA constructs with specific parts (promoters, coding sequences, uORFs, ERN recognition sites) and transfection ratios. These specifications were generated entirely by the biocompiler with no manual tuning.

We built all three circuits, transfected them into HEK293 cells, and collected fluorescence data (Figure 6e). In each case the cells reproduced their target pattern. The model’s prediction for each built circuit (Figure 6d) agrees with the measurement to a gRMSE of 0.076, 0.054 and 0.057 for the M, I and T pattern respectively. While the measured responses are not perfect (two of the patterns show a ~15% shift along the x axis) the target pattern remains clearly visible in each experiment.

## Discussion

Traditionally, arriving at a new cellular input-output function requires long cycles of iterative, hand-driven design. Here we close that loop in a single pass: the three target patterns went from specification to validated circuits without subsequent experimental optimization or manual changes to the generated designs.

At the core of the biocompiler software and the biomorphic networks that power it is a choice of abstraction level: we separate recipe-to-model translation from quantitative learning, specifying only known structure and letting data resolve what remains uncertain [43]. A recipe determines the circuit graph; each process is learned as a shared neural block. Three of 81 conditions in a uORF tuning matrix, together with basic single-uORF measurements, suffice to predict the rest (Figure 3). Each new experiment teaches reusable process behavior that will benefit future predictions and designs, as shown by the BMN’s ability to predict held-out topologies. Prediction errors are comparable to differences between biological repeats, although accuracy depends on topology (Figure 5c). Shared blocks also support transfer across cellular contexts: a model trained only on HEK293 predicts the tested circuits in ARPE-19 and CHO about as well as in HEK293.

In the field of synthetic biology, automated genetic design has already been demonstrated for Boolean logic [23] and RNA toehold switches [44], and analog computation has precedent in bacterial transcription-factor circuits [45], including gradient-based optimization of multi-layer networks [35]. In this work, we bring such a powerful generative design paradigm to circuits that compute graded input-output patterns, in mammalian cells.

Building beyond biological neural-like computation in DNA strand-displacement systems [32, 33] and single-layer mammalian comparators [37], we demonstrate multi-layer intracellular networks in mammalian cells, composable from a small set of orthogonal parts.

Our machine learning contribution, BMNs, connects modular neural architectures[46] with graph-structured learning[47]. Here, genetic recipes assemble process-specific blocks that share parameters across circuits and are conditioned on learned part embeddings. Whole-circuit measurements train these reusable processes, which can then be recomposed for prediction and inverse design.

BMNs can also be understood as a focused kind of cell model, bridging the gap between two more general traditions. One is mechanistic: comprehensive whole-cell models that integrate molecular processes, notably demonstrated for a minimal genome [48]. The other is data-driven: the push toward virtual cells, which starts from natural cell states measured at genome scale [49] and asks what will happen after a perturbation [50]. Here the question runs in both directions: predicting what behavior a given genetic recipe will produce, and inferring what recipe would produce a desired behavior. The learned model is therefore not only a predictor but a substrate for generative design.

A highly complementary line of work maps the genetic design space directly and at enormous scale, using high-throughput library construction and sequencing to characterize circuit variants and train a model to predict untested ones [51]. This mirrors an established approach for regulatory DNA, where massively parallel reporter assays have trained models that not only predict enhancer activity from sequence but also generate synthetic enhancers with specified activities [52]. Where such a library can be built, it provides direct, empirical coverage of that region of design space. However, the space of multi-layer compositions still grows far faster than any library can cover. The biocompiler addresses this compositional challenge because its blocks are shared and recomposable. Measurements of a handful of architectures support predictions for ones absent from training, and the same model can be searched in reverse, from target behavior to design. The two approaches could easily be combined: high-throughput characterization could sharpen the shared blocks and expand the part embedding spaces the biocompiler searches over.

Several directions follow naturally. Design currently operates over supplied scaffold architectures, and fully open-ended generation will require extending the recipe-to-graph grammar. This grammar is currently hand-authored from established circuit biology, and learning its rules directly from data is an appealing longer-term goal, alongside advances in learned neural architectures [53, 54]. Global coupling effects such as resource competition and ribosome loading are currently captured only implicitly [55, 56, 57]. Theoretical studies show that shared resources can alter the responses and decision boundaries of sequestration-based classifiers [58]; representing such coupling explicitly in the BMN could strengthen extrapolation. On the biological side, extending these transiently transfected circuits to stable programs, endogenous inputs and other neuro-morphic building blocks is another important step. The framework is indeed not specific to endoribonucleases: recipe-to-graph compilation, shared-block learning, latent encoding of parts and gradient-based inverse design could extend to other regulatory primitives whose interactions can be specified at the graph level. Other reaction mechanisms, including catalytic degradation and competitive binding, have also been modeled as perceptrons with tunable thresholds [59].

A broader opportunity for this work is to allow biological engineering to leverage experimental data in a cumulative way. Each experiment can provide knowledge about processes that recur across designs, circuits, and contexts, allowing each measured circuit to help construct new ones. As synthetic biology moves toward multicellular programs, developmental circuits and tissue-scale engineering, this ability to learn at the right level of abstraction across compositions and context becomes increasingly important. The native language of cells is analog and compositional; to program them, our tools must be too.

## Methods

### Plasmid Design and Assembly

All genetic parts and plasmids were assembled using hierarchical Golden Gate Assembly as described in previous publications [60]. First, DNA fragments or plasmids containing individual genetic parts were inserted into pL0 plasmid backbones using Golden Gate Assembly. pL0 plasmids were constructed for insulators, promoters, 5’ UTRs, genes, 3’ UTRs, and terminators. Each pL0 plasmid delivered its genetic component to a particular position in a second round of Golden Gate Assembly to construct a full transcription unit contained in a pL1 plasmid. In some cases, a further round of Golden Gate Assembly was used to combine two pL1 plasmids into a pL2 plasmid that contained both transcription units. In other cases, Golden Gate Assembly was used to incorporate pL0 parts directly into pL2 plasmids.

### Cell Culture

The poly-transfection protocol used in this publication is described in greater detail, with specific reagents, materials, and suppliers at [61]. In poly-transfection each DNA input is delivered in a separate transfection complex, so individual cells take up uncorrelated amounts of each input and a single population spans the full space of input copy-number combinations, with each cell’s input levels read out by co-transfected fluorescent markers. This is what allows a single experiment to map an entire multi-input dose-response surface, displayed as the heatmaps throughout this work: in each map the axes are the per-cell abundances of the node’s molecular inputs and the colour is its analog output (surviving output-mRNA fluorescence), rendered as a smoothed surface over the single-cell measurements (see Computational Methods, Visualization). Experiments were conducted using human embryonic kidney (HEK293) cells, except for the cell-type generalization experiments, which additionally used ARPE-19 and CHO cells. Cell cultures were kept incubated at 37°C, 5% CO_2_ and supplied with Dulbecco’s modified eagle medium (DMEM) with 10% fetal bovine serum (FBS) and 1% non-essential amino acids. Additionally, 1% Penicillin-Streptomycin-l-Glutamine (Corning) was added to cell cultures.

### Transfections

Transfections were performed as described in the referenced protocol. Cells were seeded at a density of 43,000 cells/cm^2^ in 24 well plates 24 hours before transfection. Opti-MEM was used with Lipofectamine 3000 transfection reagent and P3000 enhancer reagent as directed. Flow cytometry was performed 60 hours after transfection across all experiments unless otherwise noted.

### Flow Cytometry

Fluorescence data was recorded on BD LSR Fortessa flow cytometers. SPHERO Ultra Rainbow Calibration 9 Peaks Lot No. AJ01 Catalog NO.: URCP-100-2H were used for downstream data analysis as described in the section Data Management.

### Data Management

The contents and ratios of each genetic network element and transfection complex can be found with experimental data organized by architecture in the corresponding section of the SI. In addition, the following optional metadata items were recorded for each experiment: Transfection operator, flow cytometry operator, flow cytometer, cell line, transfection reagent, plate type, incubation conditions, transfection protocol (e.g. reverse or forward transfection), and spectral beads used. Flow cytometry data was collected using FACSDiva version 8.0.1. To screen for living single cells, gating was performed manually using side scatter and forward scatter channels. Fluorescent color unmixing was performed using calibrie, an in-lab custom python tool created by Jean Disset based on the TASBE workflow [62]. All fluorescence values were converted to mean equivalent fluorescence (MEF) units of EBFP2.

### Plotting

Custom perceptually uniform colormaps were used in all heatmaps (see Computational Methods).

### Data calibration and preprocessing

Raw fluorescence data from flow cytometry were processed using Calibrie, a Python pipeline for quantitative fluorescence calibration inspired by the TASBE protocol [63] and CytoFlow [62]. Processing steps include: spectral unmixing via weighted least-squares linear compensation from single-protein controls; conversion to molecules of equivalent fluorophore (MEF) using Spherotech calibration beads (URCP-100-2H), with bead peak assignment via optimal transport; and mapping of all protein abundances to a common reference scale (EBFP2). Living single cells were gated using forward and side scatter channels. All fluorescence values reported in this work are in “Pacblue” MEF units of EBFP2. Calibrie is available at https://github.com/jdisset/calibrie.

### Recipe-to-compute-graph compilation

Each genetic design is specified as a *recipe*: a declarative description of DNA constructs (transcription units with promoter, uORF, coding sequence and ERN recognition site slots) and co-transfection ratios. The biocompiler turns each recipe into a circuit-specific compute graph by first constructing a graph representing biological information flow (DNA→RNA→Protein edges with part identities as edge metadata), then applying declarative graph-rewriting rules that encode known biological interactions. For example: merging identical DNA sources, creating ratio-weighted aggregation nodes for co-transfection, and generating ERN inhibition nodes by matching ERN proteins to their cognate RNA recognition sites. Rules are applied iteratively until the graph reaches a stable state. The output is a directed acyclic graph where each node has a known process type and the circuit architecture is determined entirely by the recipe and the rules, with no learned components.

### Neural blocks and parameter sharing

Each node in the compute graph is assigned a *neural block*: a small multi-layer perceptron (MLP) that predicts the quantitative input-output behavior of its corresponding biological process. All nodes of the same type across all circuits share the same MLP weights, conditioned on part-specific embedding vectors. This weight sharing provides implicit supervision: each new circuit constrains the shared blocks from a different compositional context, which means the system of functional equations becomes increasingly determined as the training corpus grows.

Discrete biological parts (promoters, uORFs, ERN types) are mapped to continuous vector embeddings. Embeddings are parameterized as variational distributions (mean and log-standard-deviation), with reparameterized sampling during training and KL regularization against a unit Gaussian prior. This variational formulation forces the latent space to be smooth and centered: neighboring parts occupy neighboring positions, enabling continuous traversal by a gradient-based optimizer during design. The straight-through estimator (STE) allows gradients to flow through the discrete quantization step, so that the forward pass can use discrete part assignments while the backward pass treats quantization as identity.

### Training procedure

The biocompiler is trained on flow cytometry data from multiple circuit designs simultaneously. Each experiment produces a distribution of single-cell fluorescence measurements throughout the input space, not a single value, and predictions are matched to individual measurements rather than to per-condition summary statistics.

For the regression results reported in this paper, the training objective combines three terms: a per-sample mean squared error between predicted and observed single-cell outputs, with per-output weighting and a soft split between coupled (graph-internal dependent) and independent outputs; an inverse-pair consistency term that, for each forward/inverse process-block pair automatically discovered in the graph (source, transcription, translation and output), penalizes the residual of a forward-then-inverse cycle through the blocks with sampled rate embeddings; and a KL regularization of the variational part embeddings against a unit Gaussian prior, which keeps the latent space smooth and centered. The inverse-consistency and KL weights are ramped during training. The same loss function supports additional terms for distributional prediction (sorted-output MSE, quantile pinball, CDF calibration) that are disabled in the regression configuration used here.

Optimization uses AdamW [64] with global gradient norm clipping and a warmup-cosine learning-rate schedule. The training set is expanded progressively for the predictions shown in this study: models for each circuit class include all simpler circuit types, so that each prediction tests generalization to an unseen circuit architecture. Configurations and training scripts for every run reported in this paper are released alongside the code.

Model names record which topology classes were in the training set: S single-ERN, M multi-ERN single-layer, Lc cascade, Lbl and Lbr the two bandpass orientations, N no ERN. A model is therefore named for what it has seen.

**Table 1.** Trained model used for each reported prediction.

| Figure / section | Model | Classes in training |
| --- | --- | --- |
| Figure 3 | building_blocks | elementary circuits only |
| Figure 4, regions | SN | single-ERN, no-ERN |
| Figure 4, cascades | SMN | + multi-ERN single-layer |
| Figure 4, bandpass BL-TR | SMLcLbrN | all but left bandpasses |
| Figure 4, bandpass TL-BR | SMLcN | all but both bandpasses |
| Figure 5 | SMLcLbILbrN | all classes (learned blocks, not a held-out prediction) |

### Generative design

In design mode, the trained blocks and part embeddings of the BMN are frozen and the biocompiler solves the inverse problem: given a target input-output pattern, find discrete part assignments and transfection ratios that produce the desired behavior. Gradient descent through the frozen forward model operates on continuous part embeddings and transfection ratios, minimizing the distance between predicted output and target. Simultaneously, transcription unit (TU) masking via the Hard Concrete distribution with an L0 penalty selects which circuit elements to retain. Optimization proceeds in three phases: exploration (all TUs active, low sparsity pressure), pruning (increasing L0 weight), and commit (hard-prune remaining TUs). A two-timescale learning rate separates TU mask updates from embedding optimization.

The design loss combines Sinkhorn optimal transport distance, local normalized cross-correlation (LNCC), and scale-invariant MSE, with penalties for ratio bounds and TU count. Because the variational embeddings enforce a smooth latent space, gradient descent traverses a continuous landscape even though the final output must be a discrete part assignment. After optimization, continuous embeddings are quantized to the nearest valid discrete part via masked nearest-neighbor lookup using the STE, and the resulting circuit is verified by forward prediction. The final graph is then bio-compiled to a human-readable recipe and DNA construct list for experimental implementation, by applying in reverse the same graph-rewriting rules used in the forward direction.

### Visualization

All heatmaps in this work display smoothed dose-response maps generated by fixed-bandwidth Gaussian kernel regression (Nadaraya-Watson smoothing) rather than binned histograms. We evaluate this estimator by scatter-splatting instead of gathering neighbors at each grid point: every single-cell measurement is deposited onto a regular high-resolution lattice by cloud-in-cell interpolation, accumulating a few moment buffers (local counts and kernel-weighted sums of the outputs); the buffers are then blurred once with a separable Gaussian, and the value at each grid cell is recovered as the ratio of the blurred buffers. This returns exactly the same fixed-bandwidth Nadaraya-Watson estimate as a per-pixel neighbor average, but its cost scales with the number of cells and the grid size instead of a nearest-neighbor query at every grid point, which keeps high-resolution maps and 3D volumes tractable. The result is a smooth, continuous surface without the discretization artifacts of binned heatmaps. Grid cells with fewer than 20 measurements within the kernel radius are masked as no-data, and all heatmaps use custom colormaps with perceptual uniformity.

### Prediction accuracy metrics

The models in this paper are regression models: at each location in the input space they predict the average single-cell output. We therefore measure error at the level of the dose-response surface rather than of individual cells. For each dataset we fit a smooth response surface to the measurements with the same fixed-bandwidth Gaussian kernel smoother (Nadaraya-Watson) used for the heatmaps, and evaluate it on a regular grid spanning the input space, in rescaled fluorescence units (nominally 0–1). Outputs are rescaled with a symmetric-log transform anchored so that the working range of the assay maps onto the unit interval. The transform is not clipped, so a surface that runs above that range takes values above 1.

Our error metric, the **grid RMSE (gRMSE)**, is the root-mean-square difference between two such surfaces over that grid. Because both surfaces are smoothed, single-cell scatter is averaged out and does not enter the comparison, so the gRMSE reports disagreement in the underlying input-output behavior. When the two surfaces are a model prediction and the matching experiment, we call it the **pred gRMSE**. When a circuit was measured in independent experiments (biological replicates), the same quantity computed between two repeats and averaged over repeat pairs is the **rep gRMSE**: it says how much a circuit’s behavior changes when the experiment is repeated. Both are on the same scale and directly comparable, and a model whose pred gRMSE on a circuit is at or below that circuit’s rep gRMSE predicts it about as well as repeating the experiment would.

### Software and hardware

The biocompiler is implemented in Python using JAX [65] for differentiable computation and automatic vectorization, with Optax [66] for optimization. Flow cytometry calibration (raw fluorescence to standardized MEF units, spectral unmixing, automatic and interactive gating, and quality metrics) uses Calibrie (https://github.com/jdisset/calibrie). Configuration, CLI generation and pipeline composition use Dracon [67] (https://github.com/jdisset/dracon), a YAML-based configuration system developed alongside this work to manage the layered training, prediction, plotting and design experiment configurations. All figures in this work, from genetic-circuit and network schematics to the dose-response maps and analysis panels, are produced with JeanPlot (https://github.com/jdisset/jeanplot), a standalone composable plotting library built on top of Matplotlib; the biocompiler-specific panels are layered on top of it in biocomp-tools (https://github.com/jdisset/biocomp-tools and biocomp). Parallel execution and monitoring of multi-condition training and design sweeps use Broodmon, a parallel command runner and process orchestrator built on top of Dracon. Training and design were performed on a single workstation with an NVIDIA RTX 5090 GPU and 128GB of RAM.

### Code availability

The biocompiler source code is available under the MIT license at https://github.com/jdisset/biocomp (core framework) and https://github.com/jdisset/biocomp-tools (accompanying tools). Calibrie is available at https://github.com/jdisset/calibrie, Dracon at https://github.com/jdisset/dracon, and jeanplot at https://github.com/jdisset/jeanplot. The Broodmon parallel runner, and all scripts, jobs and configuration files used to train the models, plot experimental results, generate predictions and design circuits in this work, will be released under the same license prior to publication, with versioned snapshots archived on Zenodo.

## Data availability

The calibrated single-cell flow cytometry measurements and the trained biomorphic neural network models that support the findings of this study will be deposited on Zenodo prior to publication.

## Code availability

The biocompiler framework, including the biomorphic neural network training and inverse-design pipelines, together with the Dracon configuration system, the Calibrie flow cytometry calibration library and the plotting utilities used to produce all figures, is available as open source on GitHub under the MIT license; the repositories are listed in the Code availability subsection of the Methods. The Broodmon parallel experiment runner and the scripts, jobs and configuration files specific to this study will be released under the same license prior to publication, with versioned snapshots archived on Zenodo.

## Acknowledgments

We thank the members of the Weiss lab for helpful discussions and feedback throughout the project. In particular we thank Gustavo Aguilar and Yiming Wan for all the valuable feedback on the manuscript and figures, and João Costa for his generous help with the computing hardware used in this work. We thank Christian Cuba Samaniego for his contributions during the early ideation of the neuromorphic circuits presented here [26].

This work was supported by the National Science Foundation through PROGENIC (Programmable Organoid Intelligence using Neuronal Networks implemented by Gene Circuits), CyberOrganoids (CPS: Medium: Microrobotics-enabled differentiation control loops for cyber-physical organoid formation), Fine-grain generation of multiscale patterns in programmable organoids using micro-robots (GCR: Collaborative Research), and Evolvable Living Computing: Understanding and Quantifying Synthetic Biological Systems’ Applicability, Performance and Limits (Collaborative Research); by the Defense Advanced Research Projects Agency (DARPA) through BioSPARC (Biological Synthesis through Perceptron-based Artificial Regulatory Circuits), Synthetic Biology Inspired Machine Learning, and BIOSYNC (Bioelectronics for the Delivery of Synthetic Therapeutics with Wireless Control); by the National Institutes of Health through Genetically Programmed Pancreatic Organoids with Self-Adaptive Multi-Lineage Population Control, Synthetic gene sensors and effectors to redirect organoid development, and Programmed Differentiation Circuits for Organoids using Meso-Microfluidics; and by the Air Force Office of Scientific Research through A synthetic biology programming language and foundational control theory with application to guided multicellular mammalian 3D shape shifters. Additional support came from NIH grant R21 CA257980 and US Army Research Office grant W911NF2120109. G.K.A.W. was supported by an Imperial College London Schrödinger scholarship.

## Author Contributions

J.D. conceived the biomorphic neural network framework, in which biological circuits are represented as graph compositions of shared neural blocks; developed the theoretical formulation and architecture; implemented the biocompiler; and designed, performed and interpreted all computational experiments of training, ablation, prediction and inverse design. J.D. analyzed and curated all experimental data collected by C.V.D.M., developed all supporting computational infrastructure used in the study (including the flow-cytometry calibration software and the configuration, parallel-execution and plotting tools described in Methods), contributed to the design and selection of the genetic circuits used for training and the showcase experiments, wrote the majority of the manuscript, and produced most of the main and supplementary figures.

C.V.D.M. performed all wet-lab experimental work, including building the genetic circuits, designing and running the poly-co-transfection experiments, and collecting and calibrating the flow cytometry data. C.V.D.M. was the principal contributor to the manual design and selection of the genetic circuits used in the training sets and in the network topologies showcased in this study, making the practical design decisions throughout the experimental campaign. C.V.D.M. contributed substantially to manuscript writing (especially earlier versions) and fully authored the experimental Methods sections.

G.K.A.W. contributed to the design and selection of genetic circuits used in training and the showcase experiments, contributed to manuscript writing, data plotting, and to earlier versions of several main-text and supplementary figures.

A.M. and W.X. contributed their expertise in neuromorphic programming to the preparation and review of the manuscript.

E.P.T. and C.B. contributed their expertise in computational modelling and machine learning to the preparation and review of the manuscript.

A.B.M. assisted with the implementation of early components of the biocompiler codebase.

S.-H.K. assisted with a subset of the cell-line experiments.

R.W. supervised all aspects of the project, guided the research direction and the interpretation of results, contributed to the design and selection of the genetic circuits used in training and the showcase experiments, secured funding, and contributed extensively to the writing and editing of the manuscript.

J.D. and C.V.D.M. contributed equally to this work.

## Competing interests

J.D., G.K.A.W. and R.W. are co-founders in an early-stage company founded to develop technologies related to this work. The remaining authors declare no competing interests.

## Additional information

**Correspondence and requests for materials** should be addressed to Ron Weiss.

## Notes

### Summary of Updates

Updated some citations and wording about prior art.

